# Kidney-specific lipid metabolism underlies stable chemical individuality in domestic cats and other felids

**DOI:** 10.1101/2025.09.01.673589

**Authors:** Shota Ichizawa, Jana Caspers, Reiko Uenoyama, Mizuki Morisasa, Naoko Goto-Inoue, Tamako Miyazaki, Yusuke Suzuki, Nozomi Nakanishi, Yasuyuki Endo, Masako Izawa, Tetsuro Yamashita, Stefan Schulz, Masao Miyazaki

**Affiliations:** Department of Bioresources Science, The United Graduate School of Agricultural Sciences, Iwate University, 3-18-8 Ueda, Morioka, Iwate 020-8550, Japan; Institute of Organic Chemistry, Technische Universität Braunschweig, Hagenring 30, 38106 Braunschweig, Germany; Department of Marine Science and Resources, College of Bioresource Sciences, Nihon University, Kanagawa, Japan; Cooperative Department of Veterinary Medicine, Faculty of Agriculture, Iwate University, Iwate 020-8550, Japan; Department of Materials and Applied Chemistry, College of Science and Technology, Nihon University, Building No.2, 1-5-1 Kanda Surugadai, Chiyoda, Tokyo, 101-0062, Japan; Kitakyushu Museum of Natural History & Human History, 2-4-1 Higashida, Yahatahigashi-ku, Kitakyushu, Fukuoka 805-0071, Japan; Joint Faculty of Veterinary Medicine, Kagoshima University, 1-21-24 Korimoto, Kagoshima, Kagoshima 890-0065, Japan

## Abstract

Domestic cats uniquely accumulate lipid droplets (LDs) in renal proximal tubular epithelial cells, but their chemical composition and biological functions remain unknown. Here, we identify a distinctive renal specialization in felids: lipid droplets enriched in branched-chain fatty acids (BFAs). These BFAs accumulate as triglyceride components and are excreted in urine as free fatty acids with stable and individually distinctive profiles. Behavioral assays show that cats discriminate urine samples based on BFA composition, even against complex volatile backgrounds, suggesting BFAs as robust, metabolically encoded olfactory signatures for individual recognition. Comparative analyses of non-domesticated felids reveal that BFA production is broadly conserved but exhibits both species-specific and individual variation. These findings demonstrate how a core metabolic pathway has been evolutionarily co-opted into a chemical signature system. This discovery highlights an unusual integration of renal physiology with social communication and expand our understanding of lipid metabolism, organ-level adaptation, and the evolution of chemical individuality in mammals.

**Significance statement:** Unique lipid droplets in felid kidneys reveal how organ-specific metabolism can be adapted for social communication.

## Introduction

Across vertebrates, core organs such as the heart, lungs, liver, kidneys, spleen, and gastrointestinal tract exhibit remarkably conserved architectures and functions, reflecting deeply shared developmental programs and essential physiological roles (*1*). Yet, as external morphology adapts to habitat and lifestyle, the internal structure and cellular organization of these organs can also diverge under lineage-specific selective pressures, generating functional innovations while retaining fundamental constraints (*2, 3*).

A striking example occurs in healthy domestic cats (*Felis silvestris catus*), whose kidneys are unusually yellow due to the abundant, persistent accumulation of lipid droplets (LDs) in proximal tubular epithelial cells (Fig. 1A, (*4, 5*)). In mammals, physiological LD accumulation in internal organs is uncommon and typically limited to specialized contexts: hepatocytes transiently store triglycerides after dietary intake (*6*), enterocytes accumulate LDs during nutrient absorption (*7*), and steroidogenic cells in the adrenal cortex, ovaries, and testicular interstitium maintain LDs as cholesterol reservoirs for hormone synthesis (*8*). Such accumulation is usually transient, nutritionally regulated, or linked to endocrine functions requiring rapid lipid mobilization. By contrast, chronic LD accumulation in organs, such as in hepatic steatosis, is generally pathological (*9, 10*). The feline renal cortex thus poses a physiological paradox: a non-steroidogenic, non-absorptive tissue that consistently contains abundant LDs under normal conditions. The ubiquity of this trait among individuals suggests a deeply rooted evolutionary adaptation in domestic cats, and possibly across the Felidae family.

**Fig. 1.**
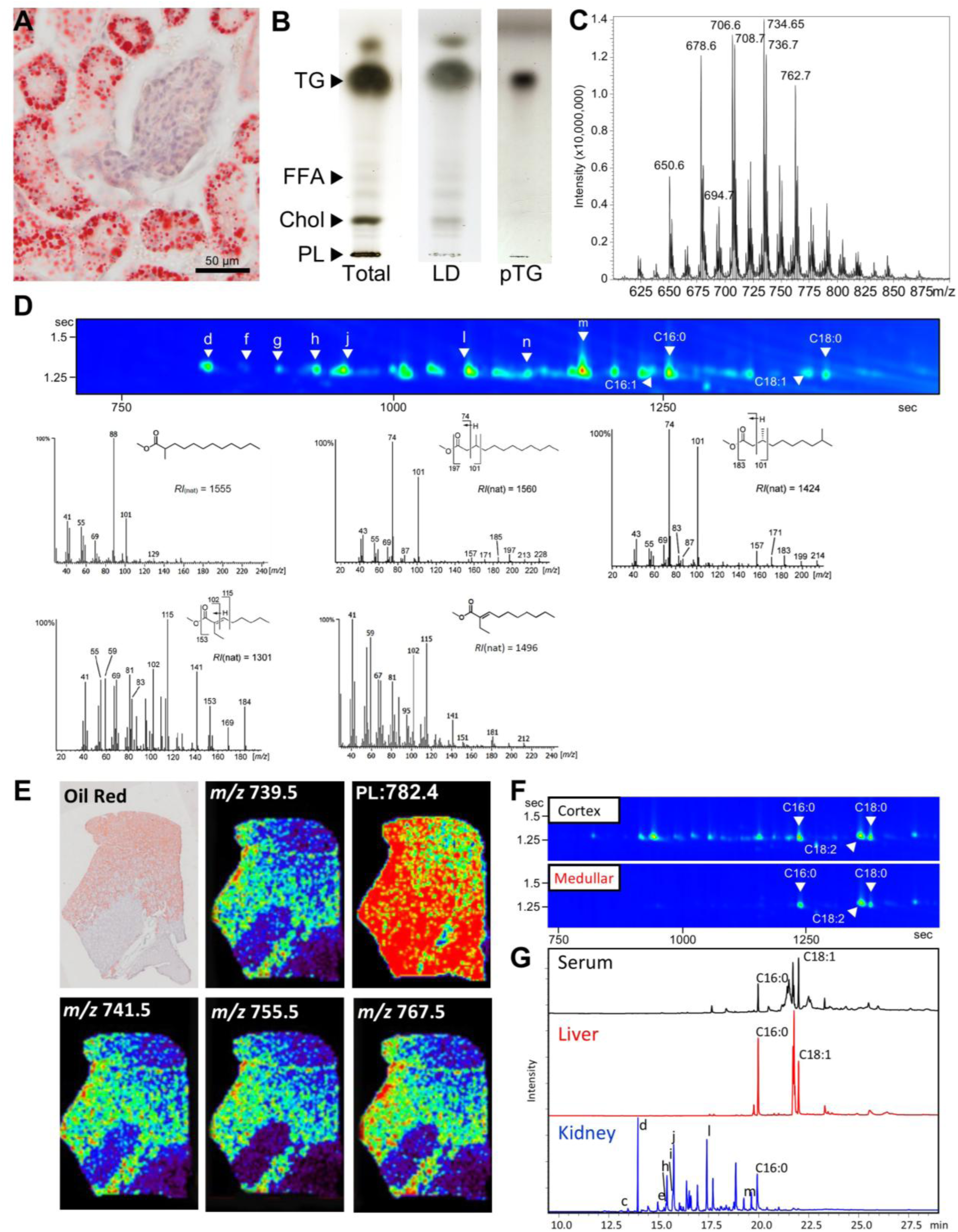
Histological and biochemical analyses of renal lipid droplets in cats. **A.** Frozen kidney section from a male cat stained with Oil Red O, showing lipid droplets (LDs) in tubular epithelial cells. Scale bar = 50 μm. **B.** Thin-layer chromatography (TLC) of renal lipid samples. TLC plates were stained with 3% copper acetate in 8% phosphoric acid to visualize lipid classes in total kidney lipids (Total), LDs isolated from kidneys (LD), and triglycerides purified from LDs (pTG). Each spot corresponds to lipids extracted from 2 mg of kidney tissue (wet weight). Major lipid classes are indicated: triglycerides (TG), free fatty acids (FFA), cholesterol (Chol), and phospholipids (PL). **C.** Mass spectrum of triglycerides purified from renal LDs, analyzed by LC/ESI-MS. Monoisotopic ions corresponding to individual triglyceride species are indicated. **D.** GC×GC/MS analysis of triglyceride-derived BFAs. Counter plots of total ion chromatograms (TIC). Representative EI mass spectra of methyl ester derivatives prepared from triglycerides purified from renal LDs. Letters correspond to individual BFAs listed in Table 2. **E.** MS imaging of feline kidney sections, showing phospholipids (PC 34:1) and BFA-containing triglycerides in sequential sections. BFAs were highly co-localized with Oil Red O–positive LDs. **F–G.** Tissue distribution of BFAs in cats. **F.** Counter plots of GC×GC/MS TIC of methyl ester derivatives of triglycerides from cortex and medulla regions of cat kidneys. **G.** GC-MS TIC of methyl ester derivatives of total lipids from feline serum, liver, and kidney.

Although this kidney phenotype has been described histologically for over a century (*11, 12*), both the chemical composition and physiological function of these LDs have remained unknown. Histological surveys have also documented renal LDs in several non-domesticated felids, including leopard, jaguar, lion, cheetah, serval, fishing cat, and tiger (*13*), but their biochemical composition has never been determined. Here, we identify a distinctive lipid class in feline renal LDs, medium-chain branched-chain fatty acids (BFAs), previously unreported in mammals. We show that BFAs are excreted into urine as free fatty acids (FFAs) with stable and individually distinctive profiles, and that cats can discriminate among individuals based on these profiles. Comparative analyses reveal that BFA production is conserved across non-domesticated and domestic felids but exhibits both species-specific and individual variation. These findings uncover a previously undescribed renal specialization, demonstrating how a basic metabolic pathway can be evolutionarily co-opted for chemical communication, and expand our understanding of lipid metabolism, scent signaling, organ-level adaptation in mammals, and the evolution of chemical communication.

## Results

### Identification of Unusual Branched-Chain Fatty Acids from Feline Kidneys

To determine chemical profiles of LDs, we isolated lipid droplets (LDs) from renal cortex homogenates by a simple gradient centrifugation method. Thin-layer chromatography (TLC) compared chemical components between the isolated LDs and total lipids extracted from the renal cortex using chloroform/methanol (2:1) (Fig. 1B). Triglycerides were predominant components in both preparations, where FFAs and cholesterol were present at lower levels in the LD fraction than the total lipid. Phospholipids were notably absent from the LDs, indicating that feline renal LDs are largely composed of triglycerides with minimal membrane-associated lipids.

Triglycerides were further purified from the LDs by normal-phase liquid chromatography (Fig. 1B) and then analyzed by electrospray ionization (ESI) mass spectrometry (MS). Several ammonium-adducted precursor ions were detected, including *m/z* = 650.6, 678.6, 694.7, 706.6, 708.7, 722.7, 734.7, 736.7, and 762.7 (Fig. 1C). MS/MS analyses of these ions revealed neutral losses of fatty acids (FAs) corresponding to individual acyl chains, although the chemical structures of these FAs could not be fully resolved by ESI-MS alone. To identify them, we performed two-dimensional gas chromatography (GC×GC)–MS of methyl ester derivatives prepared from the triglycerides. Based on electron impact (EI) mass spectra, gas chromatographic retention indices, and direct comparisons with synthetic standards, GC×GC– MS identified several BFAs not previously described in mammals (Fig. 1D, and Table 1). BFAs were classified into six structural types; 3-methyl, 3,9-dimethyl, 2-methyl, 2-ethyl, and 2,4-diethyl BFAs, which were predominantly medium-chain FAs (C10:0–C14:0). Together, the ESI-MS and EI-MS data indicate that most triglycerides in renal LDs contained at least one BFA, typically combined with common FAs such as oleic acid (C18:1) or palmitoleic acid (C16:1).

**Table 1.** Lists of branched fatty acids (BFAs) identified from cat urine.

| Compound | Chemical | RI<sup>*1</sup> | MW<sup>*2</sup> |
| --- | --- | --- | --- |
| a | (*E*)-2,4-Diethylhex-2-enoic acid | 1231 | 170 |
| b | (*E*)-2-Ethyl-2-octenoic acid | 1301 | 170 |
| c | 3-Methyldecanoic acid | 1362 | 186 |
| d | (*R*)-3,9-Dimethyldecanoic acid | 1424 | 200 |
| e | 2-Methylundecanoic acid | 1455 | 200 |
| f | 3-Methylundecanoic acid | 1460 | 200 |
| g | (*E*)-2-Ethyl-2-decenoic acid | 1496 | 198 |
| h | 3,10-Dimethylundecanoic acid | 1532 | 214 |
| i | 2-Methyldodecanoic acid | 1555 | 214 |
| j | 3-Methyldodecanoic acid | 1560 | 214 |
| k | 3,11-Dimethyldodecanoic acid | 1622 | 228 |
| l | (*E*)-2-Ethyl-2-dodecenoic acid | 1694 | 226 |
| n | 3-Methyltetradecanoic acid | 1758 | 242 |

<sup>\*1</sup> RI: Retention index of each BFA methyl ester is shown.
<sup>\*2</sup> MW: molecular weight of BFA

To confirm that these BFA-containing triglycerides originated from LDs in proximal tubular epithelial cells, we performed imaging MS. Sodium-adducted ions at *m/z* 782.5, corresponding to phosphatidylcholine (PC 34:1), were detectable throughout the sections (Fig. 1E). In contrast, sodium-adducted ions corresponding to BFA-containing triglycerides (*m/z* 739.6, 741.6, 755.6, 757.6, 767.6, 783.6, 795.6, and 823.6) were highly localized to the renal cortex in regions corresponding to Oil Red O-positive LDs and were nearly absent in the medulla, consistent with the paucity of LDs in that region.

We next compared the BFA content of triglyceride fractions from renal cortical and medullary LDs. GC×GC–MS analysis revealed that BFA-derived methyl esters were significantly enriched in cortical LDs relative to medullary LDs (Fig. 1F). Moreover, GC–MS analysis of total lipid extracts from serum and other organs detected BFAs exclusively in the kidneys, confirming their tissue specificity (Fig. 1G). Together, these results demonstrate that LDs in proximal tubular epithelial cells of the feline renal cortex constitute a previously unrecognized lipid class in mammals: triglycerides carrying structurally distinct BFAs, highlighting the kidney as a unique site of BFA biosynthesis and storage, revealing an unexpected renal lipid metabolism in mammals.

### Urinary Excretion of Branched Fatty Acids

Previous studies have reported the presence of large amounts of LDs in feline urine [Fig. 2A, (*14, 15*)]. To determine whether triglycerides carrying BFAs are excreted in urine, we analyzed total urinary lipids from male domestic cat samples by GC×GC–MS following treatment with trimethylsilyldiazomethane, which methylates FFAs but not triglycerides. In addition to large, late-eluting peaks corresponding to methyl esters of common C16 and C18 FAs, several BFA methyl esters were identified in cat urine (Fig. 2B). Importantly, only few BFAs were detected by GC–MS in kidney samples treated with trimethylsilyldiazomethane, suggesting that triglycerides stored in renal LDs undergo hydrolysis and are excreted as FFAs in the urine.

**Fig. 2.**
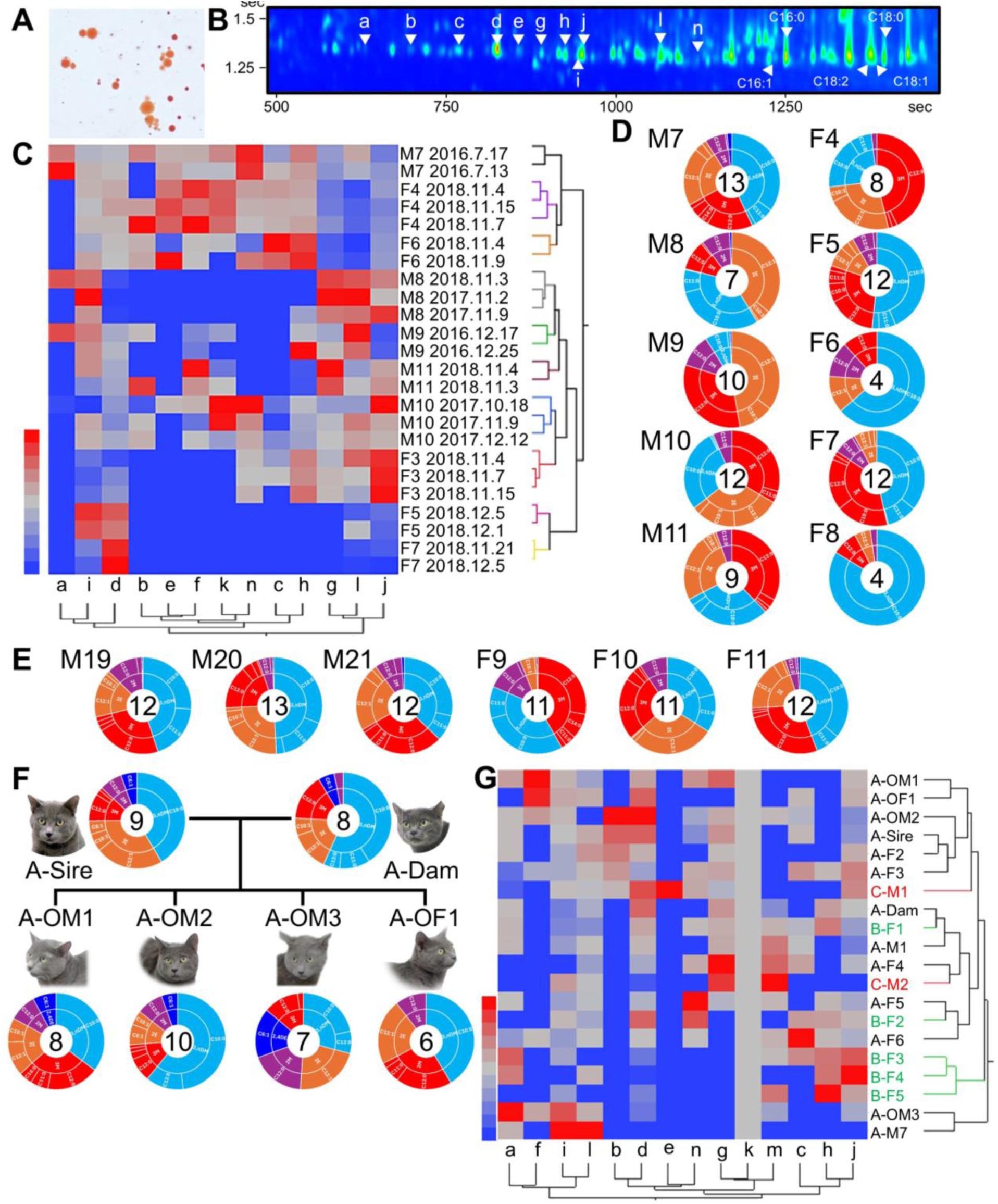
Urinary branched-chain fatty acids (BFAs) in domestic cats. **A.** Oil Red O staining of urinary lipid droplets. **B.** Counter plots of GC×GC/MS total ion chromatograms (TIC) of methyl ester derivatives of urinary total lipids. Letters correspond to BFAs listed in Table 2. **C.** Heat map and dendrogram of 13 BFA methyl esters detected in 24 urine samples collected from ten cats (five males, five females). Relative contents (%) were calculated from GC×GC/MS TIC peak areas, and Ward’s minimum variance was used for hierarchical clustering. Sampling dates are indicated; all data sets are shown in Data S2. **D.** Average urinary BFA profiles of ten cats. Pie charts show mean chemical compositions (%) of 13 BFAs based on two or three urine samples from each individual. Color codes are as follows: red, 3-methyl BFAs; cyan, 3,9-dimethyl BFAs; orange, 2-ethyl BFAs; violet, 2-methyl BFAs; blue, other 3-methyl BFAs. Fatty acids are denoted as x:y, where x indicates the carbon chain length and y indicates the number of double bonds (e.g., 10:0, saturated 10-carbon fatty acid). **E.** Renal BFA profiles of six cats (three males and three females). Pie charts show the relative compositions (%) of 13 BFAs, calculated from GC×GC/MS TIC peak areas of methyl ester derivatives of total renal lipids. **F.** Family-based analysis of urinary BFA repertoires. Urinary BFA profiles of a sire, a dam, and their four offspring (three males and one female, as in F) are shown. Each pie chart represents the relative composition of 13 BFA methyl esters. **G.** Heat map and hierarchical clustering of urinary BFA repertoires from the family shown in G together with 14 additional cats. Profiles from multiple individuals and families are compared across colonies. Cat colonies are indicated as A, B, and C (see Table SX). M and F indicate male and female, respectively. Letters correspond to individual BFAs listed in Table 2.

### Stable Individual and Familial BFA Patterns

We next assessed inter-individual variation in urinary BFAs using GC×GC–MS across 24 urine samples from five males and five females (each sampled on ≥2 days). Hierarchical cluster analysis (HCA) grouped all 24 samples into ten clusters, each corresponding to a single cat, with no clear sex-based segregation (Fig. 2C). Relative abundances of 13 BFAs were highly variable among individuals but showed similar patterns within individuals (Fig. 2D).

To test whether these individual urinary signatures reflect tissue-level variation, we analyzed renal BFAs in six cats (three males, three females). Distinct individual patterns were observed with no apparent sex-specific differences (Fig. 2E). Together, these results indicate that urinary BFA composition is individual-specific and temporally stable over both short (days) and long (months) intervals, directly reflecting the composition of renal LDs.

We further examined urinary BFA profiles in 22 domestic cats, including a family composed of both parents and four littermates. Our family-based analysis revealed that, while each offspring maintained an individual BFA signature (Fig. 2 F), urinary repertoires often showed clear familial resemblance (Fig. 2G). Three offspring shared profiles more similar to their sire, whereas one offspring (A-OM3) exhibited a markedly different profile, indicating that individuality is preserved even within a litter. Beyond family groups, clustering analysis also suggested colony-level similarity: cats from the same colony tended to show more related BFA repertoires compared with those from different colonies. These findings support the view that both genetic factors and shared population backgrounds contribute to shaping urinary BFA profiles in cats.

### Comparative Analyses of Renal Lipid Droplets and Lipid Components across the Felidae Family

To determine whether renal LDs containing triglycerides carrying BFAs are unique to domestic cats or represent a broader trait within Felidae, we examined kidney samples from multiple non-domesticated species. These included large *Panthera* species—lions (*Panthera leo*), tigers (*P. tigris*), and leopards (*P. pardus*)—as well as caracals (*Caracal caracal*), cheetahs (*Acinonyx jubatus*), pumas (*Puma concolor*), and the small Iriomote cat (*Prionailurus bengalensis iriomotensis*). Oil Red O staining revealed LDs in the renal cortex of most species, closely resembling those in domestic cats (Fig. 4A). Although many kidneys from non-domesticated felids exhibited age- or disease-related pathology, such as tubulointerstitial nephritis, LDs were consistently detected in preserved proximal tubular regions surrounding the glomeruli. Interestingly, Iriomote cats displayed markedly smaller LDs than other felid species. In a leopard, LDs were detectable in the renal collecting ducts near the pelvis. Cheetahs were the notable exception, with LDs largely absent. These observations suggest potential species-specific regulation of renal lipid storage.

To assess whether renal BFA accumulation represents a conserved biochemical trait within Felidae, we analyzed total kidney lipid extracts from frozen or formalin-fixed samples of these species using GC–MS. Several BFA methyl esters identified in domestic cats were also detected in the kidneys of non-domesticated felids, indicating a broadly shared capacity for renal BFA production (Fig. 3A). However, the BFA repertoires of non-domesticated species were generally simpler than those of domestic cats. Tigers and Iriomote cats predominantly produced 2-methyl BFAs, with 3-methyl BFAs present at lower levels. Pumas showed only 3-methyl BFAs. Notably, 3,9-dimethyl and 2-ethyl BFAs, a unique class in our analysis, were exclusively detected in domestic cats. These findings indicate that while renal BFA accumulation is broadly conserved across Felidae, but the composition and relative abundance of individual BFAs vary in a species-specific manner, reflecting potential evolutionary or metabolic divergence.

**Fig. 3.**
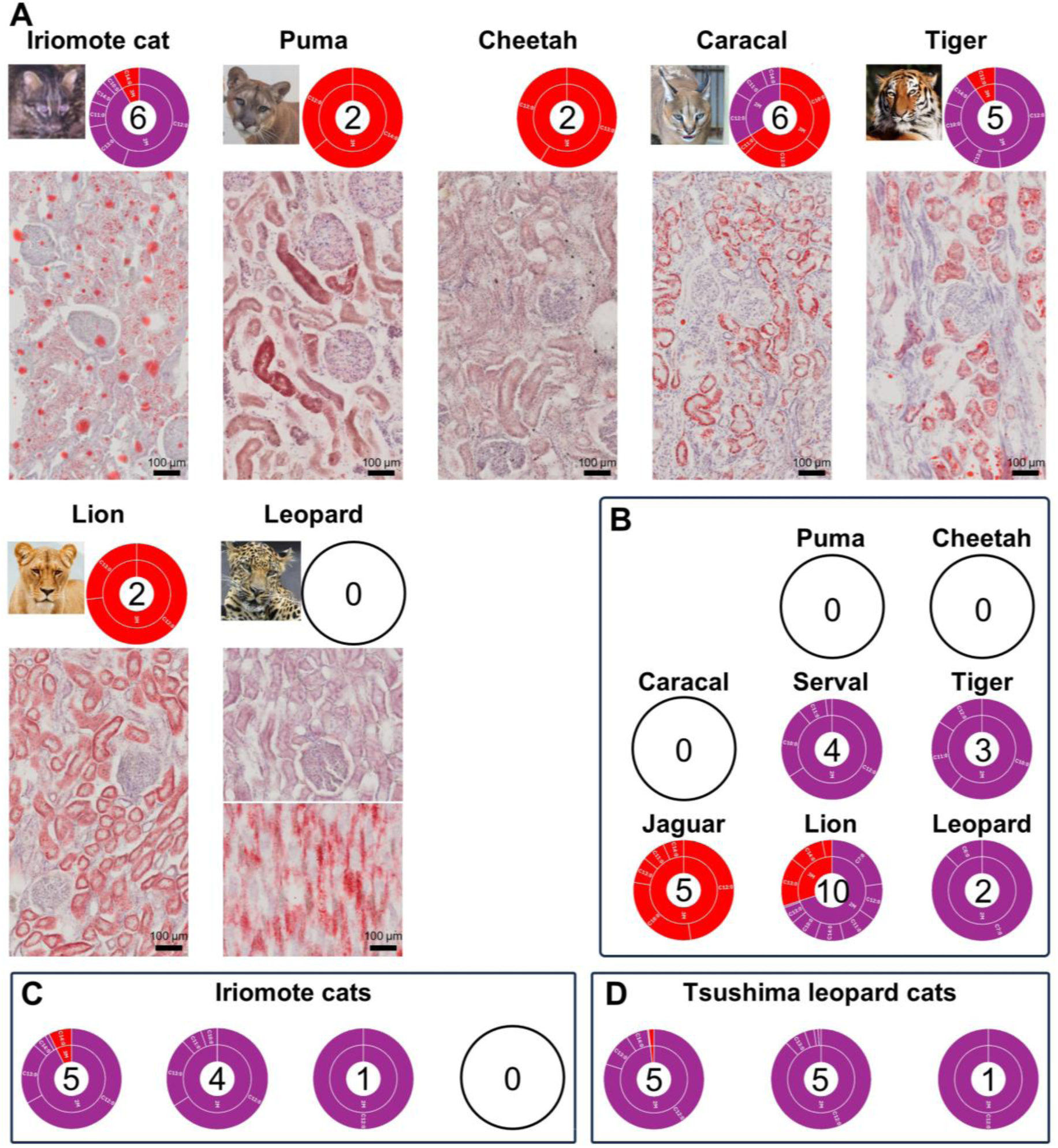
Renal and urinary branched-chain fatty acids (BFAs) in non-domesticated felids. **A.** Renal BFA compositions and histology of seven felid species. Pie charts show the relative compositions (%) of 13 BFAs in renal total lipids, calculated from GC×GC/MS TIC peak areas of methyl ester derivatives of renal total lipids. Representative kidney sections were stained with Oil Red O to visualize lipid droplets and counterstained with hematoxylin for nuclei. Scale bars = 100 μm. **B.** Urinary BFA compositions of seven non-domesticated felids. Pie charts indicate the relative abundances of 13 BFA methyl esters. **C** and **D.** Representative urinary BFA profiles of Iriomote cats (n = 26, **C**) and Tsushima leopard cats (n = 17, **D**). Four distinct repertoire patterns are shown as examples in each species. Color codes and fatty acid notations are as described in Fig. 2.

### Urinary Branched-Chain Fatty Acids in Non-Domesticated Felids

Building on our findings in domestic cats, we examined urinary BFAs in non-domesticated felids. Urine samples were collected from large and medium-sized species, including lions, tigers, jaguars, caracals, servals (*Leptailurus serval*), a puma, and a cheetah, and total urinary lipids were analyzed by GC×GC–MS after methyl esterification. BFAs were detected as FFAs in non-domesticated felids, except in cheetahs, pumas, and caracals (Fig. 3B). Most non-domesticated species exhibited simpler BFA repertoires than domestic cats, typically dominated by 2-methyl BFAs.

To assess individual variation in non-domesticated felids, we focused on two endangered small felids in Japan: the Iriomote cat (*Prionailurus bengalensis iriomotensis*) and the Tsushima leopard cat (*Prionailurus bengalensis euptilurus*). We analyzed urine from 26 Iriomote cats and 17 Tsushima leopard cats, representing a significant proportion of the ∼100 individuals estimated to remain in each population. Both subspecies predominantly excreted 2-methyl BFAs and displayed multiple individual-specific patterns (Fig. 3C and 3D). Iriomote cats showed five profile types, the most common featuring 2-methyldodecanoic acid (2-methyl C12:0) as the dominant component, with additional patterns incorporating 2-methyl C13:0 (Fig. 3C). Tsushima leopard cats exhibited four patterns (Fig. 3D), all of which were similar to the variation observed in Iriomote cats. These results demonstrate that individual-specific urinary BFA signatures also occur in non-domesticated felids, though with simpler repertoires than domestic cats.

### Olfactory Individual Signatures in Urinary BFAs

Our findings show that urinary BFAs derived from renal LDs exhibit pronounced individual specificity, suggesting that they may function as olfactory cues for individual recognition in felids. Individual recognition via scent marking underlies many key mammalian social behaviors (*16*). Conventional scent marks are composed of complex mixtures of volatile and semi-volatile compounds, yet their compositions fluctuate due to evaporation, diet, and physiological state, compromising their reliability as identity signals after deposition. This raises a central question: can renal metabolites such as semi-volatile BFAs provide sufficiently stable chemical signatures to encode individual identity?

To address this, we first examined the temporal stability of urinary BFA profiles. Cotton pads soaked with 3 mL of urine from three cats were incubated at 25 °C for 0, 3, 6, 9, 12, and 24 hours. GC×GC–MS analysis of extracted total lipids from each pad showed that the 18 samples clustered into three distinct groups corresponding to the individual cats, with minimal intraday variation compared to inter-individual differences (Fig. 4A). This suggests that urinary BFA profiles provide stable chemical signatures for at least 24 hours after scent marking.

We next conducted habituation–dishabituation assays to test whether cats could discriminate different individual BFA profiles by olfaction. Cats were first exposed repeatedly to a BFA-free urinary odor source until habituation was observed, and then to the same odor supplemented with the BFA fraction or a mixture of six authentic BFAs, which elicited significant dishabituation (Fig. 4B and 4C). Moreover, cats discriminated BFA fractions from different individuals both alone (Fig. 4D) and when mixed with the same BFA-free urinary background (Fig. 4E).

**Fig. 4.**
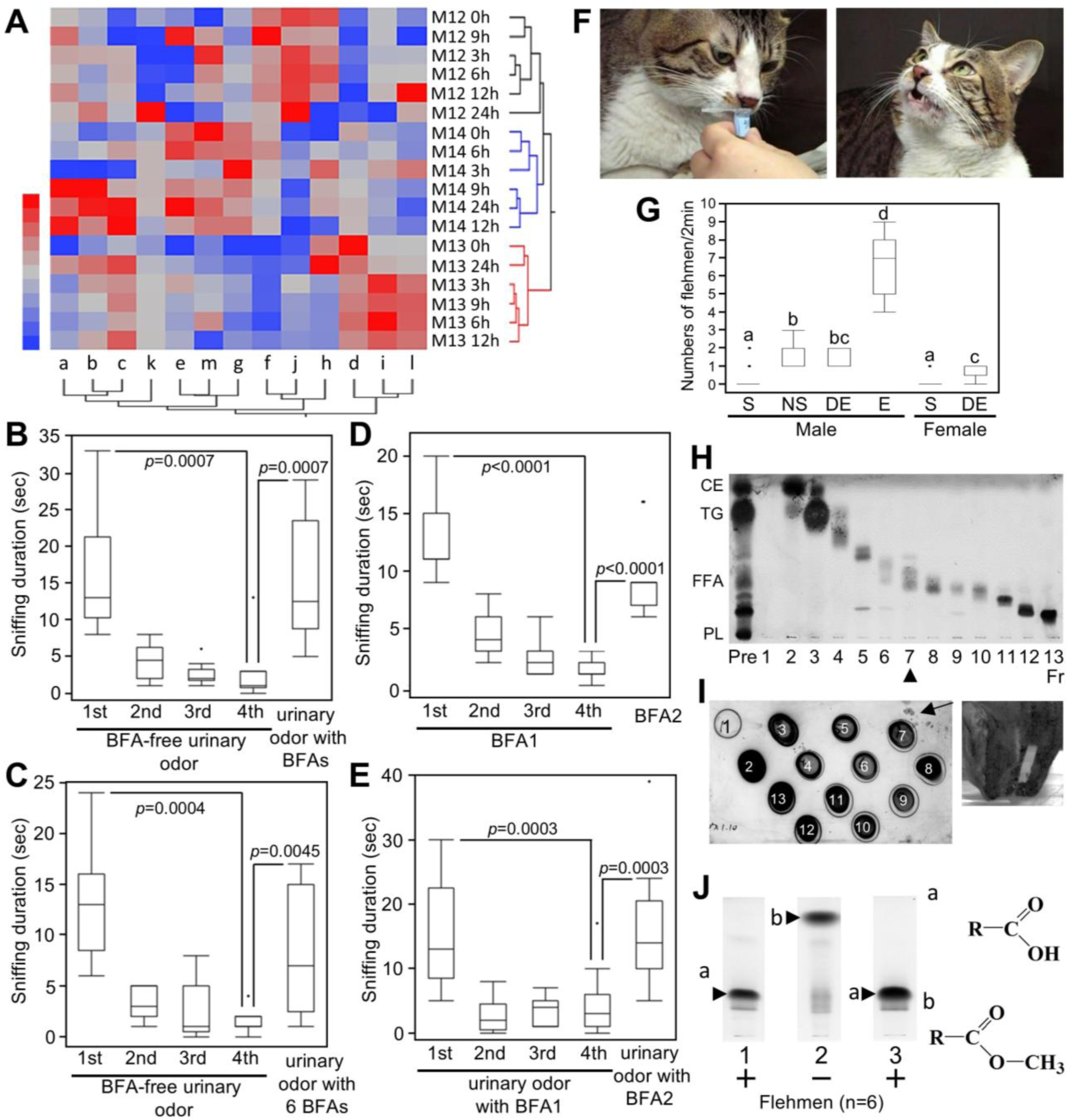
Stable urinary BFAs provide behavioral cues for individual recognition in cats. **A.** Cotton pieces that had absorbed 3 mL of urine from three male cats (M12–M14) were incubated for 0–24 h at 25 °C. GC×GC/MS was used to analyze methyl ester derivatives of total lipids extracted from the cotton pieces. The heat map and dendrogram show 13 BFA methyl esters detected across 18 urine samples (six time points × three cats). **B–D.** Habituation– dishabituation tests were conducted with five male and two female cats. In each test, the same stimulus was presented four times (60 s, 30-s intervals) in habituation trials, followed by a different stimulus once (60 s) in the dishabituation trial. Sniffing times are shown as box-and-whisker plots, and statistical differences were evaluated using Wilcoxon matched-pair signed-rank tests (p < 0.05). (**B**) Habituation: BFA-free urinary lipids; Dishabituation: the same FA-free lipids blended with the FA fraction. (**C**) Habituation: BFA-free urinary lipids; Dishabituation: the same FA-free lipids blended with six authentic BFAs (compounds a, b, c, d, i, j). (**D**) Habituation: BFA fraction from one individual; Dishabituation: BFA fraction from another individual. (**E**) Habituation: BFA-free lipids blended with the BFA fraction of one individual; Dishabituation: the same FA-free lipids blended with the FA fraction of another individual. Full data are provided in Data S6–S9. **F.** The flehmen response in a male cat. Left: sniffing a tube containing urine from another male (nasal contact). Right: the same cat displaying the flehmen response, characterized by partial mouth opening and unfocused eyes. **G.** Flehmen responses to different urine samples. Box-and-whisker plots show the number of flehmen responses in six male and three female cats after sniffing self (S) and non-self same-sex (NS) urine samples. DE and E indicate diestrus and estrous female urine, respectively. Different letters indicate significant differences among groups (p < 0.05). Detailed data are provided in Data S1. **H.** TLC of urinary lipid fractions. A stained TLC plate showing separation of 13 fractions by normal-phase HPLC. ‘Pre’ indicates total urinary lipids before fractionation, and “numbers” correspond to individual LC fractions. The triangle marks the fraction containing BFAs. **I.** Behavioral tests with LC fractions. A representative stained TLC plate spotted with 13 fractions prepared by chromatography. Such plates were used in sniffing assays with six male cats, which exhibited flehmen toward fraction 7. Arrows indicate nose imprints immediately before the flehmen response. **J.** Chemical modification of fraction 7 (the BFA fraction). Stained TLC plates of Fr. 7 before (1) and after (2, 3) treatment with trimethylsilyldiazomethane. The methyl-esterified Fr. 7 was further treated with sodium hydroxide (3) to regenerate FFAs (a) from FFA methyl esters (b). “+” and “–” indicate fractions active or inactive, respectively, in eliciting flehmen responses in six male cats.

Although these habituation–dishabituation assays demonstrated that cats can discriminate between odor sources differing in BFA content, they cannot determine whether such discrimination reflects true individual recognition. Indeed, our studies showed that cats were also able to distinguish between BFA-free urine samples from two different individuals, since their volatile compositions inevitably differ. However, these background odor profiles are generally unstable, fluctuating with time, physiology, and environment, and are therefore unreliable as markers of individual identity.

To directly test the functional role of BFAs as chemical signals, we next examined the flehmen response, a behavior in which cats raise the head and hold the mouth partially open with unfocused eyes for a few seconds after sniffing conspecific urine (Fig. 4F). Flehmen transfers urinary chemical cues to the vomeronasal organ via the nasopalatine ducts and is classically associated with males investigating estrous female urine (*17, 18*). In our assays, however, both males and females showed flehmen responses to same-sex urine from other individuals but not to their own urine (Fig. 4G). No relatedness or prior familiarity existed between subject cats and urine donors. These findings indicate that flehmen contributes to familiar/unfamiliar discrimination in cats, independent of the sexual behavior observed in intact males toward estrous female urine.

Crucially, flehmen activity was enriched in HPLC-purified BFA fractions (Fig. 4H), abolished by methylation with trimethylsilyldiazomethane, and restored after saponification, which regenerated FFAs from their methyl esters (Fig. 4I). These results indicate that BFAs are a necessary component of this response. Notably, while HPLC-purified BFA fractions consistently elicited flehmen, mixtures of six authentic BFA standards alone induced responses in only two of six cats. This suggests that BFAs act most effectively within the natural urinary odor background. The characteristic urine scent likely provides a contextual base onto which BFAs are superimposed to drive flehmen, although contributions from additional, as-yet-unidentified urinary components, including BFA species not represented in synthetic standards, cannot be excluded. In conclusion, these findings support the possibility that urinary BFAs are metabolically encoded and individually distinctive olfactory signatures in cats.

## Discussion

By identifying structurally unique BFAs from triglycerides stored in the abundant lipid droplets (LDs) of domestic cat kidneys, we provide the first biochemical characterization of a kidney-specific metabolic phenotype in felids. These BFAs are absent from other organs and serum, indicating that felid kidneys have evolved a lineage-specific lipid metabolism that biosynthesizes, stores, and excretes BFAs as urinary FFAs. Short-chain BFAs, such as isobutyric and isovaleric acids, are frequently detected in mammalian feces and anal gland secretions (*19, 20*). However, these compounds are generally produced by bacterial fermentation rather than by the host itself. Several studies have demonstrated that mammals can synthesize iso/anteiso-BFAs endogenously in specific tissues, including sebaceous glands (*21, 22*), meibomian glands (*23*), and neonatal vernix caseosa (*24*). These endogenous long-chain BFAs in mammals typically contain a single methyl branch located near the terminal end of the acyl chain. In the present study, however, we discovered that medium-chain BFAs with multiple internal methyl branches or ethyl branches accumulate as major constituents of kidney LDs in Felidae. To our knowledge, such structures have rarely been reported in vertebrates, underscoring a potentially unique metabolic adaptation associated with feline physiology.

The biological significance of excreted BFAs is likely their role as chemical signals mediating individual recognition in cat urine, given their stability and individuality. Supporting this, behavioral assays demonstrated that cats discriminate BFA profiles even against complex volatile backgrounds (*25, 26*), and chemical modification of the BFA fraction markedly reduced flehmen responses. Other urinary FFAs, such as C16:0, C16:1, and C18:1, are long-chain and nonvolatile, and thus unlikely to contribute to olfactory signaling in felids. Habituation– dishabituation assays establish BFAs as behaviorally discriminable cues, while flehmen experiments demonstrate that they also act as physiologically active stimuli engaging the vomeronasal system. In contrast to house mice, which rely on polymorphic major urinary proteins (MUPs) and associated volatile profiles for individual recognition (*27, 28*), most mammals, including felids, lack such protein-based systems. Although the habituation– dishabituation paradigm involved only limited controls and direct behavioral confirmation is impossible in cats, our results strongly support semi-volatile BFAs as the most compelling candidates for mediating individual recognition in felids, particularly when integrated into the broader mixture of urinary volatiles that produce the characteristic cat odor. Further studies, including testing other lipid classes, will be required to establish exclusivity.

The evolutionary origin of BFAs as scent signatures may parallel other vertebrate pheromones, such as androstenone in pigs (*29*) and prostaglandin F2α in goldfish (*30*), which originated as metabolic by-products later co-opted for communication. BFAs may have followed a similar route, initially serving metabolic roles before being repurposed as stable identity signals. Endogenously produced and genetically regulated, BFAs are relatively resistant to environmental noise, conferring selective advantages in territorial and social contexts. BFA signaling may have arisen from lineage-specific renal metabolism as yet unknown and been maintained by selection for reliable individual recognition.

The marked individual variation of BFA repertoires raises the question of their genetic basis in felids. We hypothesize that variation likely arises from differences in the expression of BFA biosynthetic enzymes, potentially regulated by promoter-region polymorphisms. While expression levels are usually constrained by stabilizing selection (*31*), the persistence of intraspecific diversity implies balancing selection maintaining regulatory variation under changing environmental or social conditions (*32*). This mechanism would parallel well-known systems such as ABO blood group polymorphisms (*33*) or promoter-level diversity in primate MHC genes (*34*). Our longitudinal study of a domestic cat family provides further support for heritable regulation: siblings exhibited similar BFA profiles, yet each developed a unique signature, consistent with combinatorial inheritance of regulatory variants across multiple biosynthetic genes. Such regulatory mosaicism offers a parsimonious explanation for how BFA-based chemical individuality can persist within lineages while ensuring that no two cats share an identical scent profile. While our hypothesis is based on indirect evidence from metabolic and behavioral patterns, future studies such as population-scale genomic analyses, functional validation of candidate biosynthetic enzymes, and cross-fostering or controlled breeding experiments will be required.

Comparative analyses across wild felids suggest potential species-level differences in BFA composition. Domestic cats exhibit the most diverse repertoires, including unique 3,9-dimethyl BFAs that are absent in non-domesticated felids. In contrast, wild species generally produce simpler profiles dominated by 2-methyl BFAs. One possibility is that these interspecific differences reflect evolutionary divergence in renal lipid metabolism, potentially shaped by domestication, genetic heterogeneity, and population size (*35–38*). Conversely, population bottlenecks, habitat fragmentation, and restricted breeding in non-domesticated felids (*39–42*) may have reduced genetic diversity, thereby constraining metabolic diversity and leading to simpler BFA profiles. However, sample sizes for some wild species were small, limiting statistical power and potentially underestimating intraspecific variation. These results should therefore be interpreted with caution, and it would be premature to generalize them as fixed species-specific traits. Broader sampling across populations and species will be necessary to clarify the evolutionary forces shaping BFA diversity, alongside future genomic, metabolomic, and ecological studies.

Persistent LD accumulation in non-adipose organs is rare and typically associated with lipotoxicity, as in non-alcoholic fatty liver disease or diabetic nephropathy (*43, 44*), conditions characterised by disrupted lipid homeostasis, mitochondrial dysfunction, oxidative stress, and subsequent organ damage (*45, 46*). Felid kidneys, however, tolerate triglyceride-rich LDs without pathology, suggesting protective adaptations: chronic LD accumulation in non-adipose tissues may not always be pathological. Medium-chain BFAs may be less lipotoxic than long-chain saturated fatty acids, potentially offering a natural model for safe lipid storage.

Understanding how felid kidneys manage LD accumulation and excrete BFAs may provide a useful comparative model to explore mechanisms relevant to lipid handling and toxicity. While direct clinical implications remain uncertain, these findings open an avenue for future investigation into lipid metabolism in health and disease.

In conclusion, this study elucidates the chemical structures, tissue specificity, and potential social roles of BFAs in felid kidneys. Renal LD-derived BFAs exemplify a unique physiological adaptation, demonstrating that chronic lipid droplet accumulation in non-adipose tissue can occur without pathology and may even provide a natural model for safe lipid storage. At the same time, their individual specificity and semi-volatility highlight a novel social and evolutionary function, suggesting that BFAs serve as stable chemical cues for individual recognition in felids. These findings broaden the functional scope of kidney physiology, illuminate the molecular basis of individual recognition in felids, and provide a framework for understanding how endogenous metabolites evolve into reliable social signals. Future studies to define the biosynthetic pathway for BFAs will be essential for understanding the physiological significance of this unusual, yet non-pathological, LD accumulation in Felidae kidneys.

## Materials and Methods

### Animals and biological samples

This study was conducted in accordance with local animal ethics guidelines and was approved by the Animal Research Committee of Iwate University. Thirty-six healthy, mature mixed-breed cats (1–9 years of age; 18 males, 8 females) were used for urine sample collection and behavioral assays. These cats were obtained from five feline colonies: 9 cats from colony A, 1 from colony B, 17 from colony C, 5 from colony D, and 4 from colony E. Cats were housed individually or in pairs in three-tier cages (93 cm × 63 cm × 178 cm; DCM Co., Ltd., Tokyo, Japan) under controlled environmental conditions (24 °C; 10 h light / 14 h dark cycle; lights on 07:00–17:00 h) to suppress estrous cycles in intact females. Commercial dry cat food and drinking water were provided ad libitum. Fresh urine samples free of fecal contamination were collected using metabolic cages. Sampling was attempted every 30 min between 07:00 and 19:00 h. All samples were centrifuged at 500 × g for 5 min, and supernatants were stored at −80 °C until analysis, which was conducted within one month of collection. Normal tissue samples from three male and three female cats, obtained in a previous study (*47*), were also used. These tissues had been stored at −80 °C without prior freeze–thaw cycles.

Kidney and urine samples were collected from eight non-domesticated felid species kept in zoos. Urine samples were obtained from Iriomote cats (*Prionailurus bengalensis iriomotensis*, 18 males and 8 females), Tsushima leopard cats (*P. b. euptilurus*, 10 males and 7 females), pumas (*Puma concolor*, 1 individual), cheetahs (*Acinonyx jubatus*, 1 male), caracals (*Caracal caracal*, 1 female), servals (*Leptailurus serval*, 1 male and 2 females), Amur tigers (*Panthera tigris altaica*, 1 male and 1 female), Bengal tigers (*P. tigris*, 1 male), jaguars (*Panthera onca*, 1 male and 1 female), lions (*Panthera leo*, 2 males and 3 females), and leopards (*Panthera pardus*, 2 females). Kidney tissues were obtained from Iriomote cats (2 males and 1 female), pumas (2 females), cheetahs (1 male and 1 female), caracals (1 male and 1 female), Amur tigers (1 male and 1 female), Bengal tiger (1 individual), lion (1 female), and leopard (1 male). Some kidney tissues were fixed with 4% paraformaldehyde for histological analyses, while others were stored at −80 °C for lipid extraction.

### Oil Red O staining

Lipid droplet accumulation in frozen tissue sections was visualized by Oil Red O staining. Sections mounted on MAS-coated slides (Matsunami, Osaka, Japan) were stained with 0.25% Oil Red O in 50% isopropanol (filtered) for 30 min, then counterstained with Mayer’s hematoxylin (Fujifilm Wako Pure Chemical Corp., Osaka, Japan) for 5 min. Slides were mounted in glycerol (Fujifilm Wako) and imaged using a VS120 system equipped with a VC50 camera (Evident Co., Tokyo, Japan). Images were acquired with OlyVIA v2.7 software (Evident).

Urine samples were examined for lipid droplets using Oil Red O staining solution. Lipid droplets were visualized as orange-red structures under light microscopy.

### Lipid droplet and total lipid extraction

LDs were isolated from approximately 0.2 g of frozen kidney tissue, in which the cortex and medulla were visually distinguished and separated, using a commercial Lipid Droplet Isolation Kit (Cell Biolabs, Inc., San Diego, CA, USA) according to the manufacturer’s instructions. To remove water-soluble contaminants and adjust lipid concentration, part of the LD fraction was extracted by the Bligh and Dyer method. Briefly, chloroform, methanol, and sample were mixed (1:2:0.8, v/v/v) to form a single phase. Chloroform and water (1:1, v/v) were then added, and after phase separation, the lower chloroform layer was collected for subsequent analyses.

Total lipids were extracted from frozen or formalin-fixed tissues samples (approximately 0.1-0.4 g) by the Folch method. In brief, these were incubated with 10 mL of chloroform/methanol (2/1, v/v) at room temperature overnight. Tissue residues were removed from the liquid extract by paper filtration (Whatman filter paper No. 2; Whatman International, Ltd, Maidstone, UK), and extracts were dried by rotary evaporation. Each extract was redissolved in n-hexane.

Total lipids were extracted from 1.6 mL of urine and serum samples by the Bligh and Dyer method. In brief, 1.6 mL of the urine was mixed with 2 mL of chloroform and 4 mL of methanol. After the addition of 2 mL of chloroform and 2 mL of MQ water, the aqueous and organic solvent layers were separated. The lower chloroform layer was transferred to a recovery flask and dried by a steam of nitrogen gas. The residue was redissolved in n-hexane.

### Silica gel liquid chromatography

Lipid extracts dissolved in n-hexane were applied to a silica column (8 mm i.d., packed with 7 mL Iatrobeads 6RS-8060; LSI Medience Co.) pre-equilibrated with n-hexane.

Compounds were sequentially eluted as follows: Fr. 1 with 10 mL n-hexane, Fr. 2 (for triglycerides) with 10 mL n-hexane/dimethyl ether (90:10, v/v), Fr. 3 (for BFAs) with 10 mL n-hexane/dimethyl ether (70:30, v/v), Fr. 4 with 10 mL chloroform, and Fr. 5 with 10 mL chloroform/methanol (2:1, v/v).

### Thin layer chromatography

Samples were applied to a silica gel high-performance thin-layer chromatography plate (HPTLC Silica Gel 60; Merck Millipore Co.) and developed in hexane/diethyl ether/acetic acid (70:30:1, v/v/v). Lipids were visualized by spraying with 3% copper acetate in 8% phosphoric acid followed by heating at 180 °C for ∼10 min.

### LC/MS

Lipid samples were analyzed for qualitative profiling of triglycerides using an LCMS-8040 system (Shimadzu Co., Kyoto, Japan) equipped with an ODS column (2.1 × 150mm, 5µm particle size, Inertsil ODS-3, GL Science Inc., Tokyo, Japan). Chromatographic separation was achieved with 30% solvent A (20mM Ammonium Acetate in Milli-Q water) and 70% solvent B (Isopropyl Alcohol/acetonitrile=80:20, v/v) at a total flow rate of 0.35 mL/min. The effluent was introduced into a triple-quadrupole mass spectrometer operated in positive electrospray ionization mode, in scan (*m/z* 400–1000) and product ion scan (CE 35 eV) modes, with a dwell time of 100 msec, collision gas pressure of 230 kPa, nebulizer gas at 3 L/min, and drying gas at 15 L/min heated to 250°C. Data were acquired and processed using LabSolutions v5.118 (Shimadzu Co.).

### Mass spectrometry imaging

Serial 10-μm sections were prepared using a cryostat (CM1860; Leica Microsystems, Wetzlar, Germany). Sections were mounted on MAS-coated slides (Matsunami) for Oil Red O staining, or on indium–tin–oxide (ITO)-coated glass slides (Bruker Daltonics, Germany) for MS imaging. For positive-ion mode, a matrix solution (1–2 mL of 50 mg/mL 2,5-dihydroxybenzoic acid [DHB] in methanol/water, 8:2, v/v) was uniformly sprayed onto the frozen sections using an airbrush with a 0.2-mm nozzle (Procon Boy FWA Platinum; Mr. Hobby, Tokyo, Japan). MS imaging was performed on a TOF/TOF 5800 system (AB Sciex, MA, USA) at a laser frequency of 200 Hz. The lateral resolution was set to 150 μm, and the mass range was 400–1200 m/z.

Calibration was performed using peptide standards, bradykinin and angiotensin II. Image reconstruction was carried out with Datacube Explorer software.

### Preparation of fatty acid methyl esters

Fatty acids were converted to their methyl esters by adding 0.6 mol/L trimethylsilyldiazomethane in 10% hexane (Tokyo Chemical Industry Co.) and hydrogen chloride–methanol (5 w/v%) reagent (GL Sciences Inc.) to lipid samples for methylation of FFAs and FFAs/triglycerides, respectively. Reagents were removed under a nitrogen stream, and the residues were dissolved in n-hexane for GC/MS or GC×GC/MS analyses. An aliquot equivalent to 30 µg of each tissue sample was injected for GC/MS analysis.

### GC×GC/MS

GC×GC/MS (Pegasus 4D time-of-flight MS, LECO Corp., St. Joseph, MI, USA) was used to quantify BFAs in lipid samples after methyl esterification. The primary column was a 30 m DB-5MS (0.25 mm i.d. × 0.25 μm df; Agilent Technologies, Santa Clara, CA, USA), and the secondary column was a 1.4 m Rtx-200 (0.18 mm i.d. × 0.25 μm df; Restek Corp., Bellefonte, PA, USA). The secondary column was housed in the secondary oven, and a dual-stage quad-jet thermal modulator alternately applied cold nitrogen jets (1 s each) and hot air jets (1.5 s each) to trap and refocus compounds eluting from the primary column. Helium carrier gas was supplied at 1.5 mL/min. The primary column program was 70 °C for 2 min, ramped at 8 °C/min to 270 °C, and held for 10 min. The secondary column had a +15 °C offset, with a modulation offset of 15 °C. The MS transfer line was maintained at 250 °C.

The time-of-flight MS operated with an EI source (−70 eV) over m/z 35–500, with an acquisition rate of 200 spectra/s, ion source temperature of 200 °C, and detector voltage of 1,550 V. Data were processed using LECO ChromaTOF (ver. 4.50) with chromatographic deconvolution, including TIC peak detection, alignment, and metabolite identification via NIST08 and Wiley07 libraries. As no substantial differences in TIC peak areas were observed among methyl esters of seven authentic BFAs, the relative abundance (%) of each BFA was calculated as its TIC peak area divided by the total TIC peak area of all BFAs. These relative abundances were analyzed by hierarchical clustering analysis (HCA) in JMP 12.2.0 (SAS Institute, Cary, NC, USA), which grouped samples by similarity patterns to generate a hierarchy of clusters.

### GC/MS

GC/MS (QP-2010 Ultra, Shimadzu Co.) also analyzed the chemical structures of lipid samples after methyl esterification. The GC conditions were as follows: injector temperature, 250 °C; Stabilwax column (60 m × 0.25 mm i.d. × 0.25 µm df; Restek Corp.); oven temperature program, 40 °C for 2 min, then ramped at 8 °C/min to 250 °C and held for 20 min; carrier gas, helium at 1.5 mL/min. The mass spectrometer was operated in electron impact (EI) mode at 70 eV, with an ion source temperature of 200 °C. Mass spectra were acquired in full-scan mode (m/z 35–500) at 20,000 amu/s. Peak detection from the total ion chromatogram (TIC) and tentative compound identification were performed using GCMSsolution software (ver. 4.2; Shimadzu Co.) with the NIST08 MS library. Compound structures were confirmed by analyzing authentic standards under the same GC/MS conditions and by spiking these standards into the samples. Linear gas chromatographic retention indices were calculated against a series of *n*-alkanes.

### Chemical Synthesis

BFAs were synthesized as described below. Detailed synthetic procedures, compound characterization data, and chemical identifications are provided in the Supplemental Methods.

### 3-Methylalkanoic Acids

An appropriate aldehyde was converted to methyl 2-alkenoates via a Wittig reaction.

Conjugate addition of methylmagnesium bromide under copper catalysis yielded ethyl 3-methylalkanoates, which were saponified with LiOH to produce 3-methyldecanoic, 3-methylundecanoic, and 3-methyldodecanoic acids. The enantiomers of 3-methyldecanoic acid were obtained via enantioselective conjugate addition using CuI/Tol-BINAP as ligand, affording both enantiomers in ∼90% yield and 99% enantiomeric excess.

### 3,9-Dimethyldecanoic Acid

4-Methylpentanol was converted to its bromide, and the corresponding Grignard reagent was added to methyl acrylate to produce methyl 7-methyloctanoate. Reduction with DIBAL provided 7-methyloctanal, which was directly subjected to a Wittig reaction to yield methyl 9-methyl-2-decenoate. Installation of the 3-methyl group, followed by saponification, afforded 3,9-dimethyldecanoic acid. Enantiomers were prepared by conjugate addition of methylmagnesium bromide to methyl 9-methyl-2-decenoate under chiral CuI/BINAP catalysis, achieving high yield and optical purity.

### (E)-2-Ethyloct-2-enoic Acid

In a Horner–Wadsworth–Emmons reaction, diethyl 2-methylsuccinate phosphonate was reacted with hexanal using KHMDS and 18-crown-6, yielding methyl (*E)*-2-ethyloct-2-enoate with an *E*/*Z* ratio of 2.5:1. The E-isomer was isolated by column chromatography and saponified to obtain (*E*)-2-ethyloct-2-enoic acid.

### (E)-2,4-Diethylhex-2-enoic Acid

Using a similar approach, diethyl 2-methylsuccinate phosphonate was reacted with 2-ethylbutanal under Horner–Wadsworth–Emmons conditions to give methyl (*E*)-2,4-diethylhex-2-enoate with a 5:1 E/Z ratio. The pure *E*-isomer was isolated and saponified to yield (*E*)-2,4-diethylhex-2-enoic acid.

### Habituation–dishabituation tests

Seven male cats per cage were tested. In the habituation phase, each cat received four 90 s presentations of the same stimulus at 30 s intervals to reduce sniffing time, followed by a 90 s presentation of a novel stimulus to assess dishabituation. Each cat was tested only once per day. The first experiment used two odor sources: (1) a BFA-free urinary odor prepared by mixing silica column fractions (Fr. 1, Fr. 2, Fr. 4, and Fr. 5) of total lipids obtained from a male cat not included among the subjects, and (2) the same mixture supplemented with Fr. 3 as the BFA-containing urinary odor. The second experiment used two odor sources: (1) a mixture of Fr. 1, Fr. 2, Fr. 4, and Fr. 5 as the BFA-free urinary odor source, and (2) the same mixture supplemented with six authentic BFAs (compounds a, b, c, d, i, and j). The third experiment used two BFA stimulant fractions (Fr. 3) prepared from two different male cats not included among the subjects. The fourth experiment used two odor sources: (1) a mixture of the BFA-free urinary odor source and the isolated BFA fraction (Fr. 3) from one individual, and (2) a mixture of the same BFA-free urinary odor source and the isolated BFA fraction (Fr. 3) from another individual. Each sample (equivalent to 1 mL urine) was applied to the inner surface of a 2 mL microcentrifuge tube cap, the solvent was evaporated under nitrogen, and the tube was closed immediately before testing. Trials were video-recorded (Handycam HDR-CX560 V; Sony, Tokyo, Japan), and active sniffing—defined as the nose within 1 cm of the tube opening with visible twitching—was measured with a stopwatch. Videos were independently scored by three observers (M.M., T.M., R.U.) to minimize bias. Statistical significance was determined using Wilcoxon matched-pairs signed-rank tests (JMP v12.0; SAS Institute, Cary, NC, USA).

### Behavioral assays for the flehmen response

**Experiment 1.** Flehmen responses were quantified in nine cats (six males, three females) after exposure to self and non-self urine. Assays were conducted in a test chamber (60 × 80 × 60 cm) during the light phase. Each cat was introduced ∼2 min before testing. Urine samples (0.2 mL) were presented in uncapped 1.5-mL microcentrifuge tubes, inverted three times immediately before use, and positioned ∼20 cm above the floor, corresponding to the typical height of urine spraying (Fig. 1A). Each cat was first exposed to its own urine and, after a 5-min interval, to non-self urine. Male cats were additionally exposed to diestrus urine followed by estrous urine, with a 5-min interval. Detailed information on experimental data, subjects, and urine samples is provided in Data S1. All assays were video-recorded (Handycam HDR-CX560V, Sony, Tokyo, Japan). Statistical analyses were performed using Wilcoxon matched-pairs signed-rank tests (JMP v. 12.0; SAS Institute, Cary, NC, USA).

**Experiment 2.** Total lipids were extracted from 400 mL pooled urine from six male cats using the Bligh and Dyer method. Briefly, urine was mixed with chloroform and methanol in a 3-L separating funnel (400:500:1000 mL), followed by addition of 500 mL chloroform and 500 mL water. After shaking and phase separation, the lower organic phase was collected, evaporated, dissolved in 1 mL *n*-hexane, and applied to silica gel (Iatrobeads, LSI Medicine Co.). Lipids eluted with chloroform were evaporated under nitrogen gas, dissolved in 0.4 mL *n*-hexane, and fractionated by normal-phase HPLC on a silica column (PEGASIL Silica 60-5, 4.6 mm i.d. × 250 mm; Senshu Science Co., Japan). Isocratic elution was performed with *n*-hexane/dimethyl ether (75:25, v/v) at 1.5 mL/min for 30 min. The effluent was collected into 13 fractions: Fr. 1 (0–2 min), Fr. 2–12 (30 s each), and Fr. 13 (17–30 min). GC×GC/MS confirmed that BFAs were enriched specifically in fraction 7.

For behavioral assays, each HPLC fraction (equivalent to 1 mL pooled urine) was spotted on a TLC plate at 3-cm intervals (Fig. 4I). Plates were presented to cats in the same test chamber as Experiment 1. After flehmen responses were observed, plates were collected and stained with 3% copper acetate in 8% phosphoric acid at 180 °C to visualize nose contact with flehmen active fractions.

### Preparation of fatty acid methyl esters and their saponification

To convert FFAs into their methyl esters, samples dissolved in methanol were treated with 0.6 mol/L trimethylsilyldiazomethane in 10% hexane (Tokyo Chemical Industry Co.) until the solution turned yellow. Solvents (methanol, hexane, and trimethylsilyldiazomethane) were evaporated under a nitrogen stream, and the residues were dissolved in n-hexane for flehmen assays and GC/MS. For saponification, dried methyl esters were dissolved in 2 mL chloroform/methanol (2:1, v/v), mixed with 0.6 mL of 1 M NaOH in methanol, and incubated at 40 °C for 2 h. Reactions were neutralized with 0.2 mL of 2 mol/L acetic acid and evaporated to dryness under nitrogen. The residues were dissolved in n-hexane and used for flehmen assays and TLC.

## Acknowledgments

We thank Prof. Akemi Suzuki, Dr. Minoru Suzuki, Prof. Roger A. Laine, Prof. Jane Hurst, and Dr. Tristram Wyatt for their invaluable discussions. We are also grateful to Mrs. Minoru Maita and Mr. Ryunosuke Sato for their experimental support.

## Author contributions

M.M. designed the research and wrote the manuscript with S.S. M.M., S.I., J.C., and S.S. identified chemical structures of BFAs. M.M., T.M., and R.U. performed behavioral assays. M.M., R.U., and R.S. performed GC/MS analysis. T.M. performed statistical analyses. J.C. synthesized reference compounds. All of the authors did data analysis and had discussion.

